# Functional Organization of Midget and Parasol Ganglion Cells in the Human Retina

**DOI:** 10.1101/2020.08.07.240762

**Authors:** Alexandra Kling, Alex R. Gogliettino, Nishal P. Shah, Eric G. Wu, Nora Brackbill, Alexander Sher, Alan M. Litke, Ruwan A. Silva, E.J. Chichilnisky

## Abstract

The functional organization of diverse retinal ganglion cell (RGC) types, which shapes the visual signal transmitted to the brain, has been examined in many species. The unique spatial, temporal, and chromatic properties of the numerically dominant RGC types in macaque monkey retina are presumed to most accurately model human vision. However, the functional similarity between RGCs in macaques and humans has only begun to be tested, and recent work suggests possible differences. Here, the properties of the numerically dominant human RGC types were examined using large-scale multi-electrode recordings with fine-grained visual stimulation in isolated retina, and compared to results from dozens of recordings from macaque retina using the same experimental methods and conditions. The properties of four major human RGC types -- ON-parasol, OFF-parasol, ON-midget, and OFF-midget -- closely paralleled those of the same macaque RGC types, including the spatial and temporal light sensitivity, precisely coordinated mosaic organization of receptive fields, ON-OFF asymmetries, spatial response nonlinearity, and sampling of photoreceptor inputs over space. Putative smooth monostratified cells and polyaxonal amacrine cells were also identified based on similarities to cell types previously identified in macaque retina. The results suggest that recently proposed differences between human and macaque RGCs probably reflect experimental differences, and that the macaque model provides an accurate picture of human RGC function.

## Introduction

Visual processing in the retina has been probed for decades by examining the light response properties of retinal ganglion cells (RGCs) in several animal models, including salamander, mouse, rabbit, cat, and monkey. The assembled findings indicate that although the functional organization of RGCs has some broad similarities across species -- notably, a subdivision into discrete RGC types that encode different aspects of the visual scene -- substantial anatomical and physiological differences between species suggest varying degrees of relevance for understanding human vision [1–6]. Among animal models, the macaque monkey is widely considered to be the most relevant for human vision, for several reasons: phylogenetic proximity, similar natural visual environment, similar visual behaviors, similar gross and fine structure of the retina, and similar anatomically identified RGC types [7–11]. However, few studies have compared the light response properties of human and macaque RGCs directly, and the findings have not been entirely consistent. Thus, the relevance of this important model for human vision remains incompletely understood, with consequences for scientific understanding of vision, clinical diagnosis of visual disorders, and development of technologies for treating vision loss.

Recent work has begun to probe the similarities and differences between human and macaque RGCs. One study [12] demonstrated the presence of a handful of numerically dominant functionally distinct cell types in humans, potentially consistent with anatomical results and with a distinctive feature of the macaque retina: the numerical dominance of a few cell types (see [10]). A more detailed study [13] revealed the spatial and temporal properties of four numerically dominant human RGC types that likely correspond to well-studied cell types in macaque -- ON-parasol, OFF-parasol, ON-midget and OFF-midget -- but also suggested a striking differences in temporal visual processing by ON-parasol cells, and only partially characterized their spatial properties and functional organization. A third study [14] probed responses to several kinds of stimuli and proposed a greater diversity of RGC types in human retina than macaque, emphasized by a very low reported proportion of the four major types. These apparently conflicting findings suggest the need for a fuller physiological study of the major RGC types in humans, including their spatiotemporal light response properties and functional organization, in a way that can be directly compared to a large body of work in macaque retina.

Here, large-scale multi-electrode recording from complete populations of hundreds of RGCs, combined with fine-grained visual stimulation and analysis of light response properties, were applied to isolated human retina, and the findings were compared to analysis of dozens of macaque retinas probed in the same experimental conditions. The results revealed a striking collection of quantitative similarities between the major RGC types of the two species, including spatial and temporal response properties, precise mosaic organization of receptive fields, ON-OFF asymmetries, spatial response nonlinearity, and sampling of photoreceptor inputs over space. Other cell types simultaneously recorded also exhibited properties expected from macaque retina. These results provide a foundation for further understanding human vision, and for the development of clinical interventions, based on the macaque monkey model.

## Results

To probe the functional organization of human RGCs, 512-electrode recordings [15] and visual stimulation were performed in an isolated human retina obtained from a brain dead donor, using experimental and analysis procedures developed for macaque retina [15–20]. To characterize light responses of hundreds of recorded RGCs simultaneously, fine-grained white noise visual stimulation was performed, and the spike-triggered average (STA) stimulus was computed to identify the spatiotemporal receptive field (RF) of each cell [21]. The influence of nonlinear spatial summation on light responses was also examined with recently-developed analytical methods [22]. High-density electrical images [15,23] were used to reveal the average spatiotemporal properties of the spikes produced by each cell.

### Spatiotemporal Properties of RGC Light Responses

Human RGCs exhibited spatiotemporal light response properties expected from results in macaque retina. The RF of each cell, obtained by computing the STA of high resolution white noise stimulation, had a roughly circular spatial profile, with a weaker suppressive surround (Fig. 1, top). The four major RGC types, ON and OFF parasol cells and midget cells, were identified based on clustering, mosaic organization and spatial density (see below). These cell types had biphasic light response time courses (Fig. 1, bottom) resembling those of the same cell types in macaque. Parasol cells (Fig. 1, left) had RFs roughly twice the diameter of midget cell RFs (Fig. 1, right), and more biphasic time courses, consistent with known differences in the spatial and temporal properties of these two cell types in macaque. The relative strength of the three display primaries in the response time courses were similar to those in macaque, consistent with dominant L and M cone input to these cell types. RFs of human midget and parasol RGCs were larger than their macaque counterparts at similar visual field eccentricities [13], which is unsurprising given the ∼20% larger diameter of the human eye [24,25]. In what follows, these characteristics of light response are examined in more detail for cell type identification and quantitative comparison.

**Figure 1.**
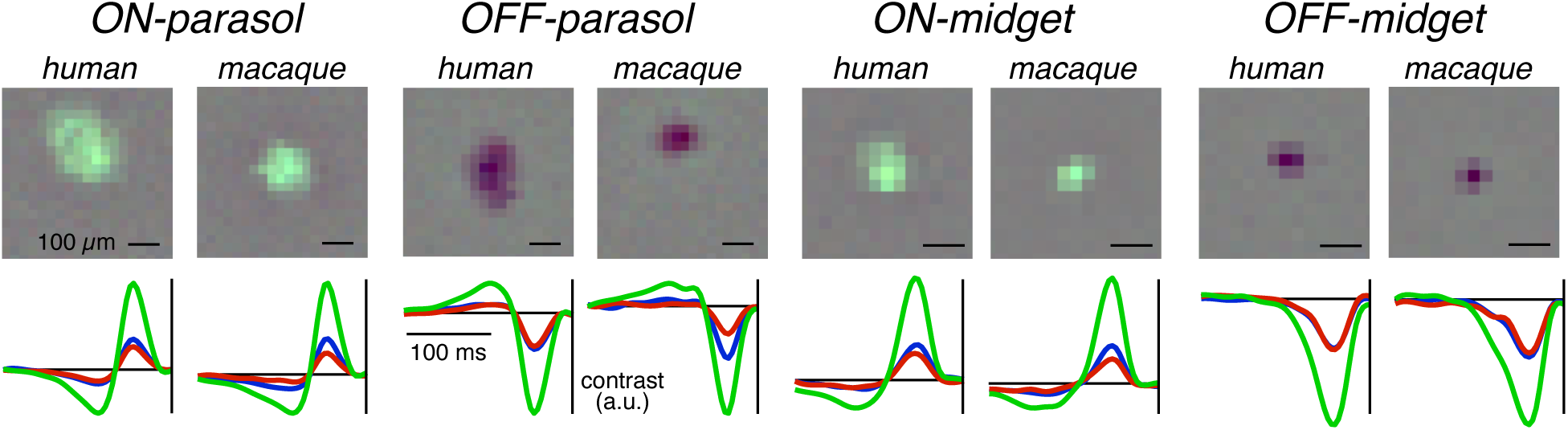
Spatial and temporal RFs of human and macaque RGCs. Each pair of panels in the top row shows the spatial RFs of a human RGC and a macaque RGC of the same type. The corresponding STA time course within the RF is shown below. Cell type classification is described below. Temporal equivalent retinal eccentricity is 12 mm in human and macaque.

### Mosaic Organization and Density of Midget and Parasol Cells

Analysis of the spatial organization of RFs confirmed the identification of distinct cell types in large-scale recordings [16,19,26]. This was accomplished by representing the spatiotemporal light responses and firing statistics of all recorded cells in a two-dimensional space using t-SNE [27], revealing discrete clusters of functionally distinct cells (Fig. 2, center). The spatial RFs of the cells in the four most numerous clusters, represented by contours of their spatial sensitivity profiles, each formed a uniform lattice, or mosaic, covering visual space (Fig. 2, flanking panels). This mosaic organization of RFs, and the stereotyped response time course in each cell type, parallels findings in other species including macaque [18,19,26], confirming that the functional clusters correspond to distinct cell types.

**Figure 2.**
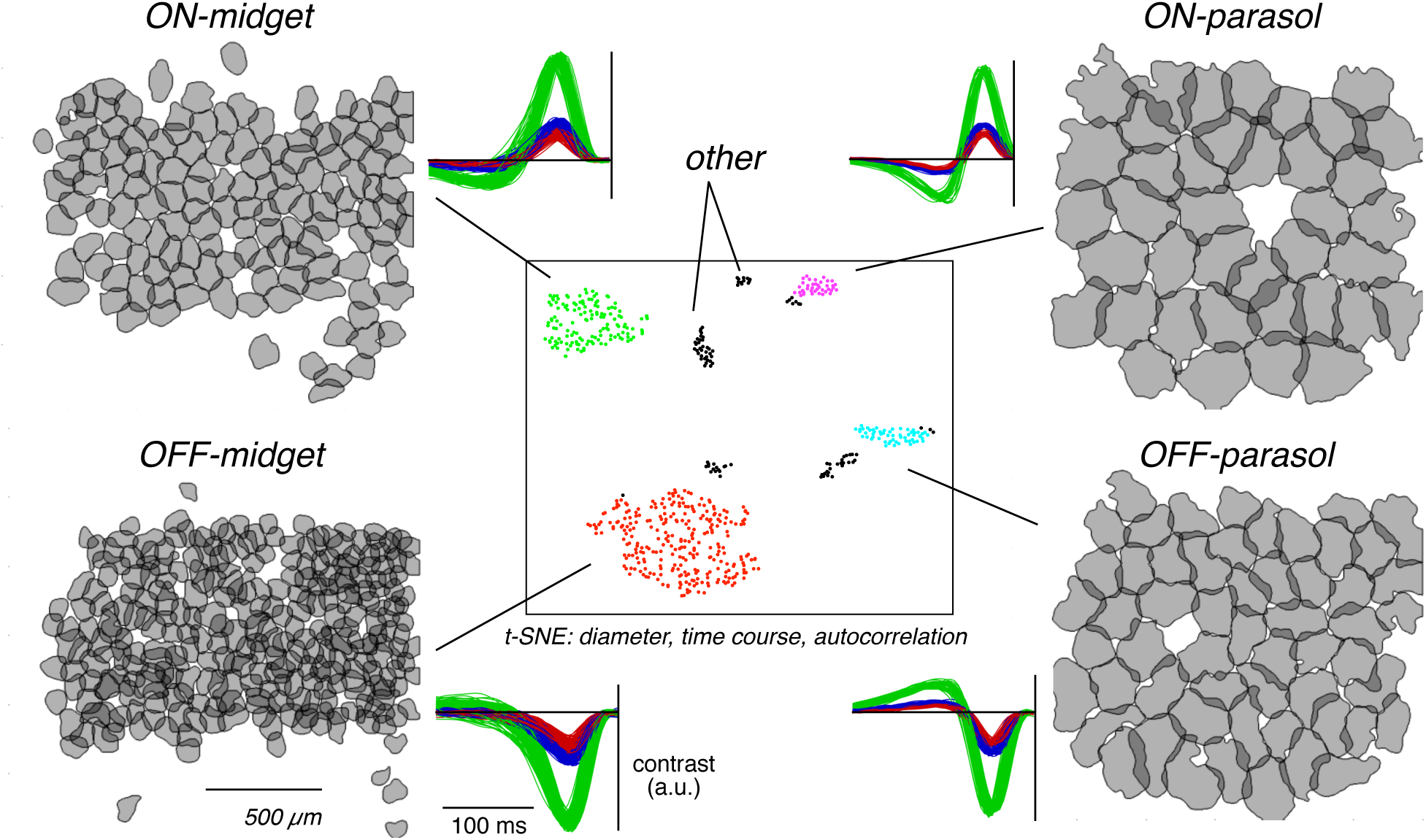
Cell type classification. Center: Two-dimensional representation of the response time courses (4 principal components), spike train autocorrelation (2 principal components) and RF area (from contour fit) for all 616 cells in a single recording (see [20]). The two-dimensional representation was obtained using t-SNE [27]. The four clusters containing the largest numbers of cells were identified and labeled with colors. Flanking panels: spatial RFs, and overlaid response time courses, of all cells from each colored cluster. Each RF is represented by a contour of the spatial profile of the STA [31]. The uniform tiling of visual space and stereotyped time courses associated with cells from each cluster confirm that the clusters correspond to specific cell types. The density and response time courses (see Fig. 1) identified the cell types as ON-parasol, OFF-parasol, ON-midget, and OFF-midget.

The identities of the four most numerous cell types were determined by analysis of their spatial densities in relation to anatomical work [16]. The spatial densities of the putative parasol and midget RGC types (Fig. 2, right and left), 59 and 360 cells/mm^2^ respectively, were comparable to the densities of the parasol and midget cells of the human retina at the same eccentricity, ∼70 and ∼200 cells/mm^2^, measured using anatomical methods [8,28]. While not in exact correspondence, these figures are broadly consistent, given the 20-40% variability in RGC density observed across human retinas [29]. Also, the variation in the ratio of parasol to midget cells, while not previously reported in the human retina, is substantial in macaque: in 20 recordings examined, the relative standard deviation of the midget to parasol density ratio was 33%. Importantly, the densities of RGC types other than parasol, midget, and small bistratified cells (which are not composed of separate ON and OFF types, and have distinctive chromatic properties not observed in the cells reported here) are expected to be at least threefold lower than the parasol cell density [30]. Also, note that polyaxonal amacrine cells have fairly high density, but are easily distinguished from RGCs based on their electrical images (Fig. 3). Thus, it is unlikely that the four observed high-density RGC types (Fig. 1,2) could be anything other than ON-parasol, OFF-parasol, ON-midget and OFF-midget. These cell type names are used in what follows.

**Figure 3.**
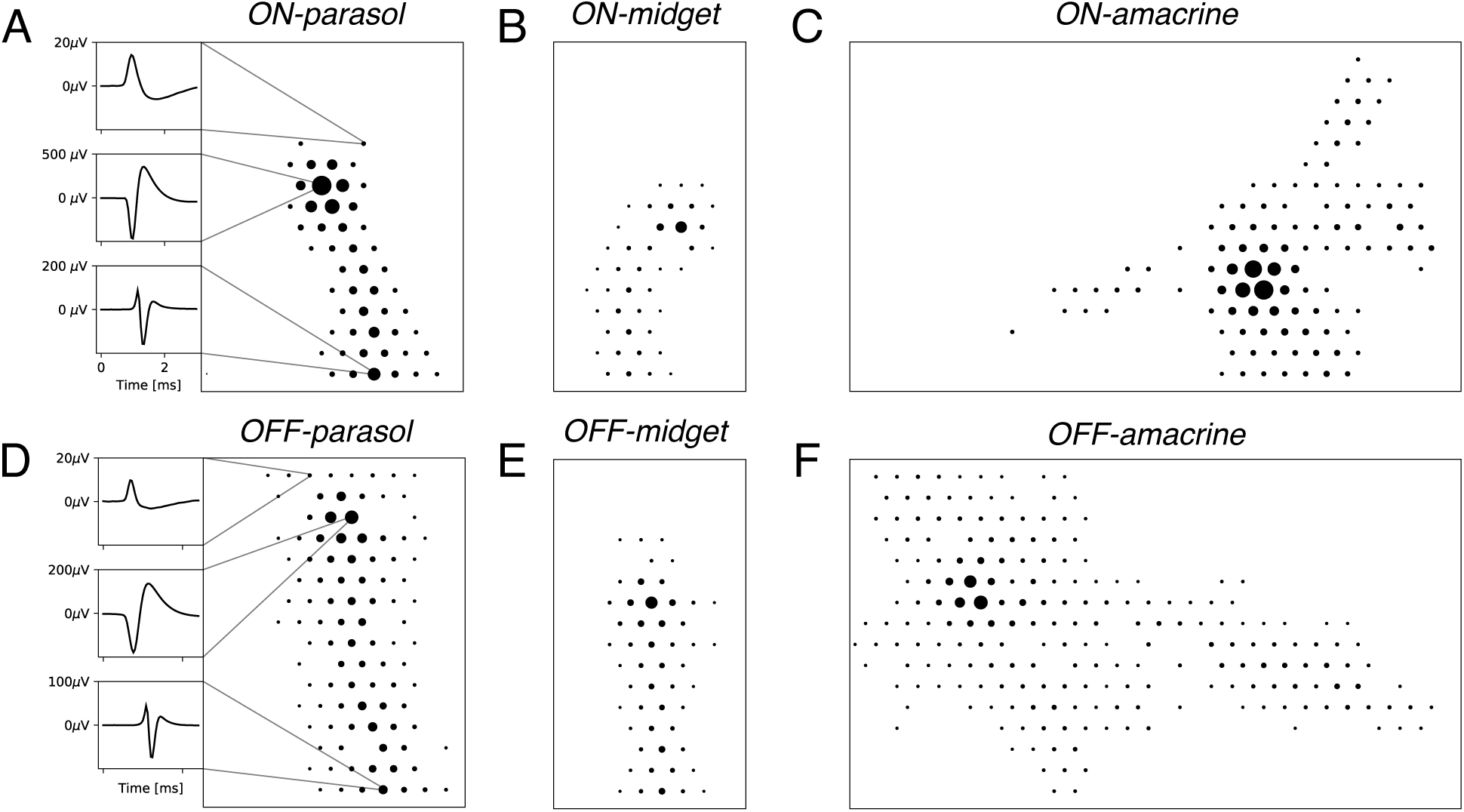
Electrical images of recorded cells. The electrical image of each cell was obtained by calculating the average spatiotemporal voltage waveform over the entire electrode array in a 3 ms time window surrounding the time of every spike from the cell ([32]; see Methods). A. The ON-parasol cell from Fig. 1. Right: over a region of the array, dot diameters show the amplitude of the recorded waveform on each electrode. Left: average voltage waveforms at each of several electrodes reveal signals recorded near the soma (middle), dendrites (top), and axon (bottom), identified by their characteristic biphasic, inverted, and triphasic shapes, respectively. The spike propagates from the soma downward along the axon. B. Spatial electrical image for the ON-midget cell from Fig. 1. C. The electrical image of an ON-amacrine cell is identified by several axons radiating outward from the soma, which is near the largest dot. D. Same as A, for the OFF-parasol cell from Fig. 1. E. Same as B, for an OFF-midget cell. F. Same as C, for an OFF-amacrine cell.

The slightly irregular RF outlines of neighboring cells of each type appeared to be coordinated, or interdigitated, interlocking like puzzle pieces in their coverage of visual space (Fig. 2). This organization resembled the results from macaque retina, in which interdigitation produced more uniform coverage of the visual field than would be produced by cells with independent spatial structure [31]. These results suggest a precise and coordinated coverage of the visual field by the major RGC types in the human retina.

The electrical image of each recorded RGC (average spatiotemporal voltage signal across the array produced by a spike) exhibited expected properties: a large biphasic spike waveform initiated near the presumed soma, opposite-sign spike waveforms at nearby dendrites presumably reflecting the current source for the somatic current sink, and propagating triphasic spikes along the axon (Fig. 3A,D, left). These electrical images closely resembled previous findings in macaque retina [32]. In addition, electrical images of other simultaneously-recorded cell types revealed spike propagation radiating along multiple axons in different directions (Fig. 3C,F). These axons are an identifying characteristic of polyaxonal amacrine cells examined previously in macaque retina [23,32].

Recent findings [13] suggest that ON-parasol cells in the human retina exhibit much more biphasic light responses than ON-parasol cells in the macaque retina, with the peak and trough of the light response time course nearly equal in amplitude. To perform this comparison in the present data, the average response time courses of human RGCs of each type were compared to assembled macaque data from 17 preparations (nine animals) obtained in the same experimental conditions (see Methods). Overall, the response time courses were similar in the two species (Fig. 4A), for all four cell types. In particular, although a small species difference was visible in ON-parasol and OFF-parasol cells, the highly biphasic time courses previously reported for human ON-parasol cells were not evident, nor was the distinction between ON-parasol and OFF-parasol cells [13] (see Discussion). For a more complete comparison, a two-dimensional space representing the response time courses was determined with t-SNE [27], using all the cells of each type from all the recordings, and each cell from each recording was represented as a point in this space. The data from human retina, though sometimes on the edge of the distribution of cells from macaque retinas, still overlapped with macaque data (Fig. 4B). Thus, the present data revealed little difference in response time course between human and macaque RGCs.

**Figure 4.**
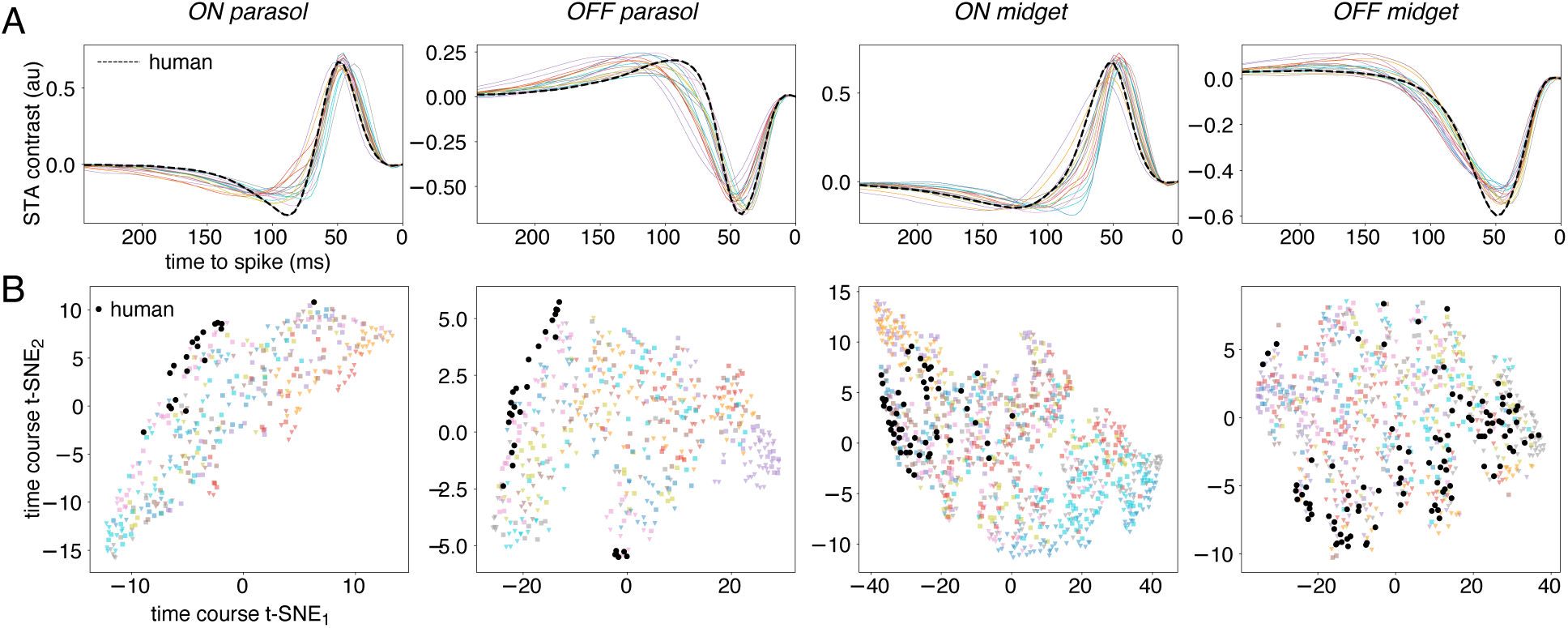
Comparison of response time courses of human and macaque RGCs. A. Average STA time course for 200 RGCs of the four major types recorded from human retina (black traces), and average time courses from 75-209 cells in each of 17 recordings of macaque retina (colored traces) using the same experimental methods and stimulus. Each time course was averaged across all cells of the same type in each preparation, and fitted with a cubic spline to interpolate between time points. B. Response time courses for every cell from the macaque and human recordings represented in a two dimensional embedding learned from t-SNE. Human cells are shown in black, cells from each macaque retina are shown in a distinct color, as in A.

### ON-OFF Asymmetries

ON and OFF cells in the human retina exhibited two asymmetries in light response properties that were consistent with previous findings [13,16,22].

First, the RFs of ON parasol and midget cells were systematically larger than their OFF counterparts (Figs. 1,2). This was quantified by fitting a two-dimensional elliptical Gaussian sensitivity profile to the spatial RF of each cell, and computing the RF diameter, given by twice the geometric mean of the standard deviation parameter of the fit along the major and minor axes. In two recordings from the human retina, the distribution of ON-parasol RF sizes was shifted to higher values relative to the distribution of OFF-parasol RF sizes (Fig. 5, left). The same was observed for ON-midget and OFF-midget cells (Fig. 5, right). Median values of the RF diameter for the four major cell types (ON-parasol, OFF-parasol, ON-midget, OFF-midget) were 144, 123, 66, and 48 µm for the first recording, and 173, 152, 64, and 46 µm for the second recording. Consistent with the RF size asymmetry and uniform spatial coverage of each cell type [26,33], the median nearest neighbor distance within each cell type was asymmetric as well, and similar to the RF diameter: 187, 143, 86, and 52 µm for the first recording, and 168, 153, 76, and 66 µm for the second recording. These ON-OFF asymmetries for parasol cells closely resembled previous observations in macaque retina [16,33], and the midget cell asymmetry resembled previous findings that are visible in published data [16,17,19,20,33] but have not been documented. Note that the present data provided inconclusive results on potential asymmetries in response time course and contrast-response nonlinearity previously reported in macaque retina (not shown; [16]).

**Figure 5.**
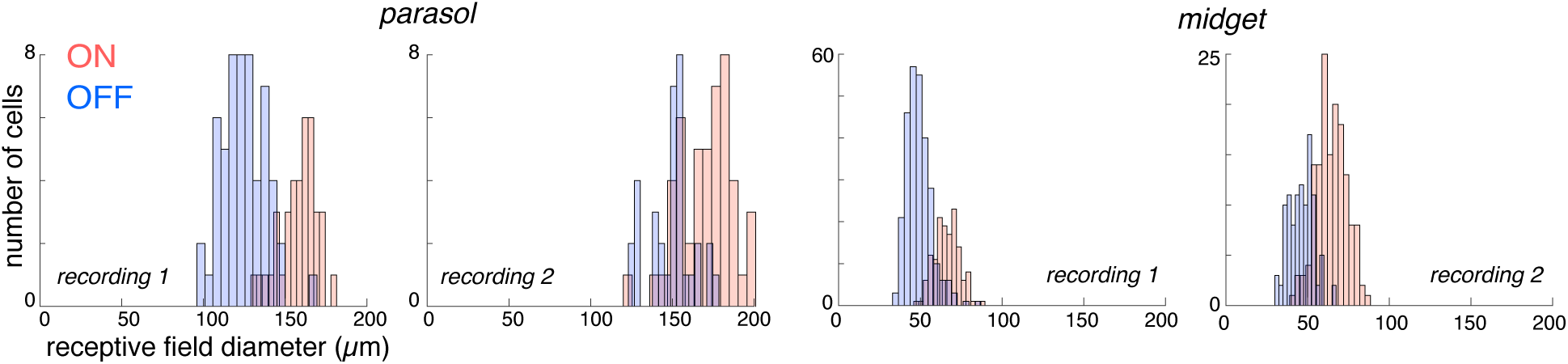
ON-OFF asymmetries in RF size. Each panel shows a histogram of RF diameters (obtained from two-dimensional elliptical Gaussian fits) for all ON and OFF parasol cells, or all ON and OFF midget cells, within a single recording from the human retina. Parasol cells are shown at left, midget cells at right; insets indicate the recording.

Second, human parasol cells exhibited nonlinear spatial summation that was asymmetric in ON and OFF cells, resembling previous findings in macaque parasol cells [22,34]. To quantify nonlinear spatial summation, a cascade model was fitted to the light responses of each cell, consisting of a first layer of several linear spatial subunits, followed by exponentiation and summation of subunit outputs, followed by a saturating nonlinearity that drives Poisson firing (Fig. 6A). Applied to individual OFF-parasol cells, fits of the model to white noise data revealed several spatially restricted subunits (Fig. 6B, right) that spanned the RF (Fig. 6B, left). The subunit model provided more accurate cross-validated prediction of light responses than the more standard and simpler linear-nonlinear-Poisson cascade model with a single spatial sensitivity profile. Specifically, as the number of subunits in the model was increased from 1, the model predicted responses more accurately, up to an asymptotic value at roughly 6 subunits (Fig. 6C, blue). However, this improvement in response prediction was smaller or absent in ON-parasol cells (Fig. 6C, red). Thus, as in the macaque retina, the nonlinear spatial summation in human parasol cells is more prominent in OFF cells than ON cells.

**Figure 6.**
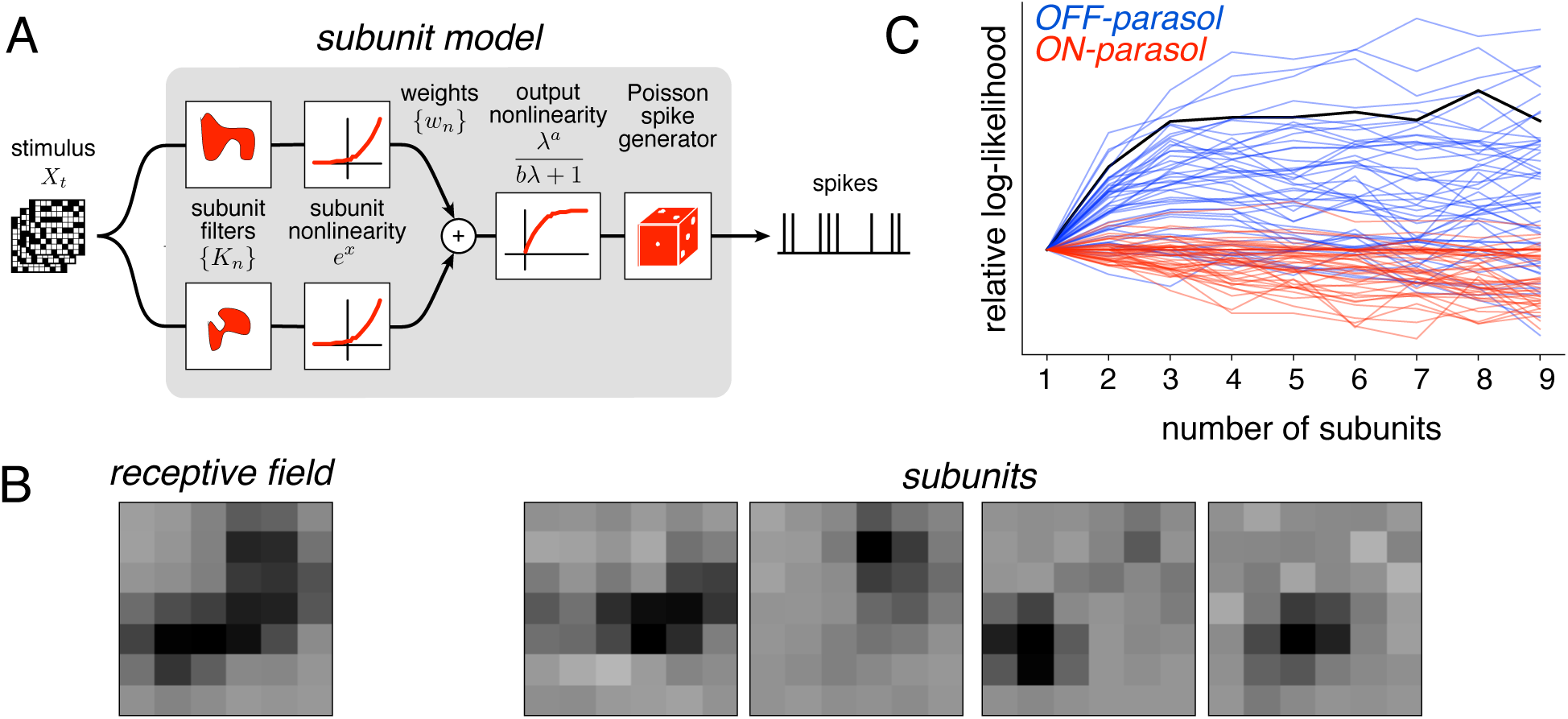
Nonlinear (subunit) spatial summation in human ON-parasol and OFF-parasol RGCs. A. Model of spatial nonlinearity, in which the outputs of multiple spatially distinct subunits are exponentiated and then summed, followed by a final saturating nonlinearity, to produce light response. A recently-developed estimation procedure efficiently estimates the subunits, weights, and nonlinearity from the responses to a white noise stimulus [22]. A special case of this model, with one subunit, is a linear-nonlinear-Poisson cascade. B. Spatial RF obtained from the STA (equivalent to a single subunit fit), and four spatial subunits obtained from a full model fit, for a single human OFF-parasol cell. C. The relative log-likelihood of the observed responses for held out data under the subunit model is shown as a function of the number of subunits, for 45 ON-parasol (red) and 58 OFF-parasol cells (blue) from the human retina, revealing that the addition of spatial subunits has a greater impact on predicting OFF-parasol responses. The performance for the OFF-parasol cell in (B) is indicated with a black trace.

### Cone Inputs to RFs

The pattern of cone photoreceptor inputs over space to the major RGC types broadly resembled results from the macaque retina. The RF of each cell, probed with high-resolution stimulation, consisted of small islands of light sensitivity spaced by ∼15 µm, and separated by regions of no light sensitivity. The spacing between these islands was similar to the cone photoreceptor spacing of ∼17 µm at the recorded eccentricity in the human retina [35]. In macaque, similar islands of light sensitivity were shown to correspond to the locations of individual cone photoreceptors [17]. The cone inputs to a given cell formed an irregular region, and were highly variable in strength over space (Fig. 7). ON-midget and OFF-midget RGCs received strong inputs primarily from L and M cones, as revealed by the relative strength of the display primaries in the response time course at the identified cone locations (not shown).

**Figure 7.**
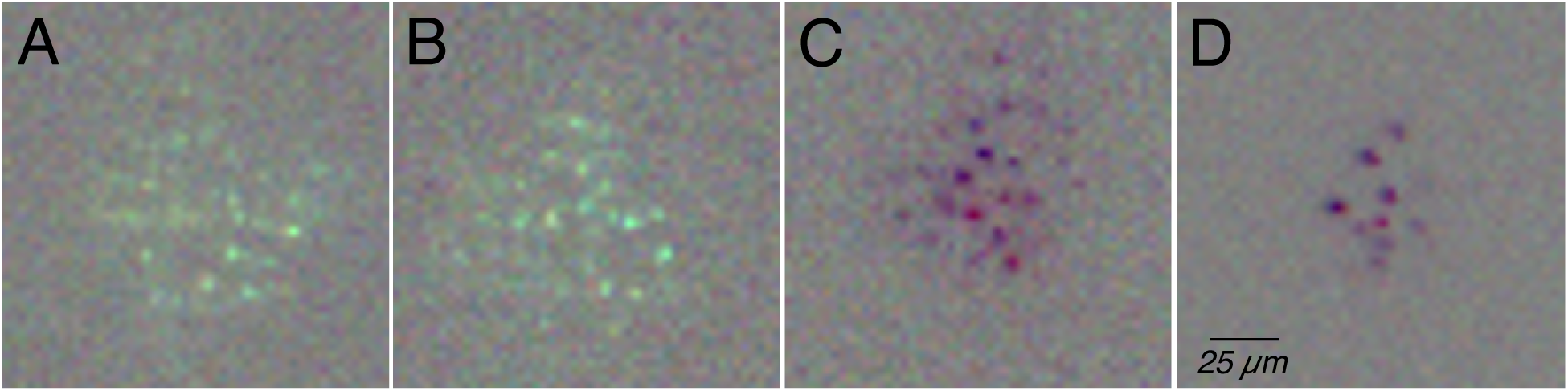
Cone photoreceptor inputs to human ON-midget and OFF-midget RGCs. A. RF of a single human ON-midget cell measured at high resolution (2.9 µm pixels), upsampled and interpolated. Isolated islands of light sensitivity, separated by regions of no light sensitivity, reveal the locations of individual cone photoreceptors within the RF [17]. The image intensity reveals the strength of the cone input, which varies considerably over space with an irregular outline. B. Same as A, for a second ON-midget cell. C. Same as A, for an OFF-midget cell. D. Same as A, for a second OFF-midget cell.

### Additional Cell Types

Additional cell types, tentatively identified from distinct clusters of light response properties (Fig. 2), also exhibited properties similar to macaque retina.

One distinct ON cell type formed a uniform mosaic (Fig. 8A). The RFs of these cells (Fig 8C,D) were larger than those of parasol cells (Fig. 1,2), but the time courses of light responses were similar (Fig. 1, Fig. 8B). These cells seemingly corresponded to the smooth monostratified cells observed in macaque retina [20]. The individual RFs exhibited a few discrete hotspots of light sensitivity (Fig 8C,D) with diameters (∼100-200 µm) much larger than the spacing between cone photoreceptors (∼17 µm) at the recorded eccentricity [35]. Spikes elicited by visual stimulation of distinct hotspots also had slightly different electrical images (not shown), an unusual property of smooth monostratified cells in macaque [20].

**Figure 8.**
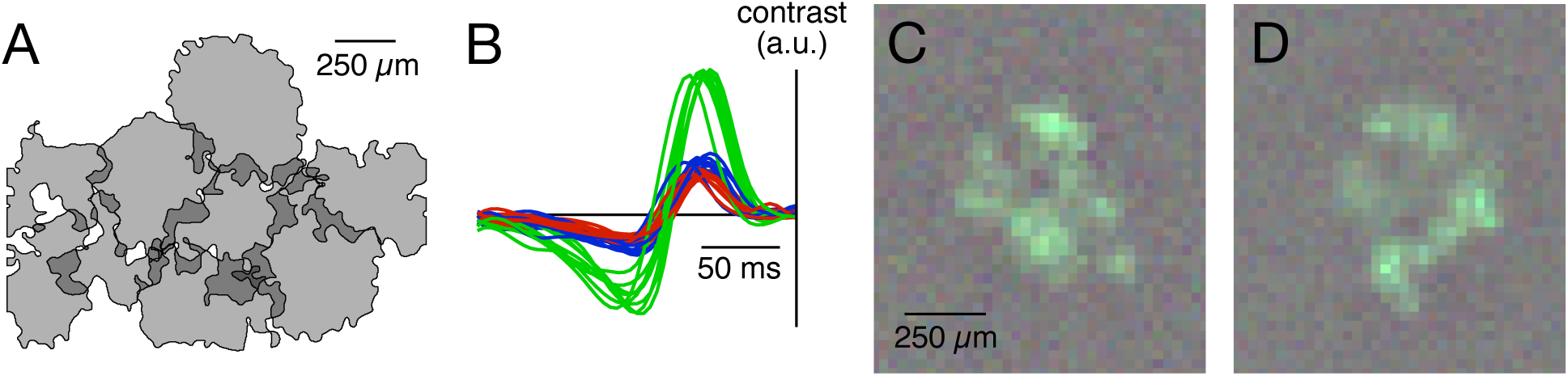
Putative human smooth monostratified RGCs. A. Contour outlines of RFs of nine ON cells of a single type (see Fig. 2, black cluster closest to ON-parasol), forming a uniform mosaic over space. B. Response time courses of the cells from A, revealing a biphasic form similar to that of parasol cells time courses (Fig. 1). C,D. RFs of two of the cells from A, with large local hot spots of light sensitivity within the RF similar to those previously observed in macaque smooth monostratified cells [20].

One OFF cell type was identified on the basis of its distinctive long, radiating electrical images as a polyaxonal amacrine cell (Fig. 3F). The RFs of these cells were characterized by irregular radiating arms of light sensitivity (Fig. 9A,C) [23]. The lack of alignment of these distinctive RF profiles with the electrically identified axons of these cells (Fig. 9B,D) suggested an origin in their dendritic structure, as expected based on the locations of their synaptic inputs and previous RF analysis [36–38].

**Figure 9.**
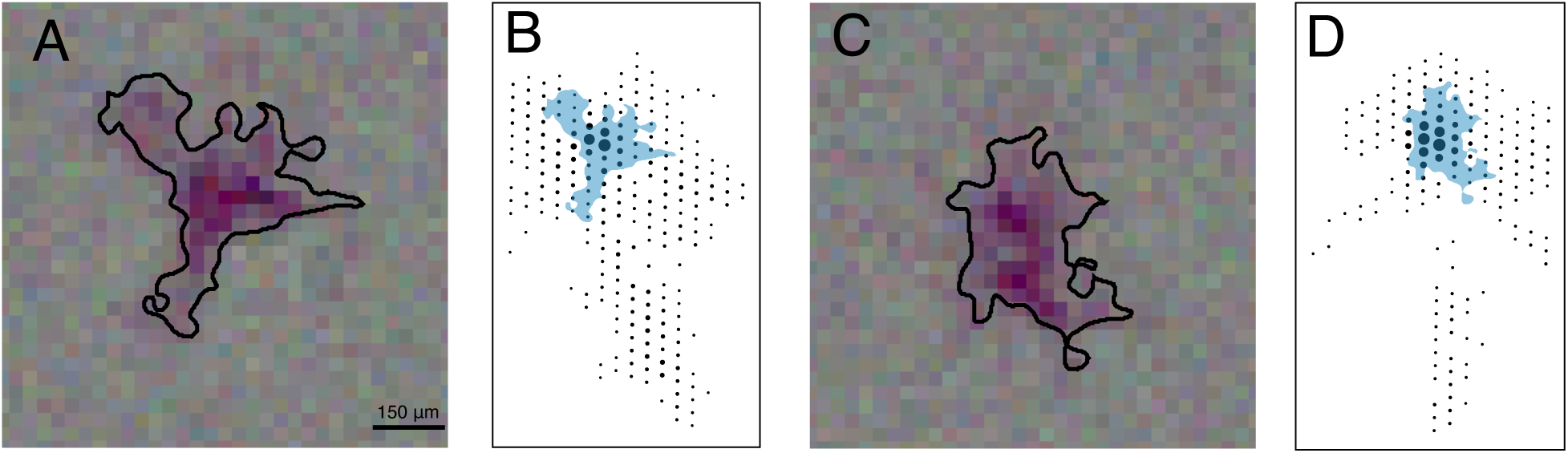
Irregular RFs of polyaxonal amacrine cells. OFF-amacrine cells exhibited distinct light responses (see Fig. 2, black cluster immediately below OFF-parasol) and unique electrical images (Fig 3C,F). A. Spatial RF of a single OFF-amacrine cell, with a contour indicating the RF outline. B. Electrical image of the same cell as A (see Fig. 3), with the RF outline overlaid in blue, revealing a lack of correspondence between the flaring features in the spatial RF (see [23]) and the extended axons in the electrical image. C. Same as A, for a second amacrine cell. D. Same as B, for the cell in C.

## Discussion

The present results reveal numerous similarities between the four numerically dominant RGC types in the human and macaque retina -- spatial and temporal response properties, precise mosaic organization of RFs, ON-OFF asymmetries, spatial nonlinearity, electrical images and sampling of photoreceptor inputs over space -- as well as distinctive features of smooth monostratified cells and polyaxonal amacrine cells. A central feature of the present results was the direct comparison to dozens of recordings from macaque retinas in the same experimental conditions. While no substantial differences were observed, the present results do not rule out the possibility of other differences between the major cell types in macaques and humans.

Although the results were obtained from a single human retina, the quantitative classification and consistent trends in hundreds of simultaneously recorded cells of different types, and the close similarity to trends observed in macaque retinas, suggest that the results will generalize. A major conclusion is that the macaque model is highly relevant for understanding the function of the human retina and visual system, particularly the aspects determined by the four major RGC types. While this has been assumed in many studies, based primarily on anatomical and behavioral findings, the present results provide a more specific and complete functional foundation.

The present results resemble those of numerous studies of major cell types in macaque retina, including many using the same techniques and experimental conditions. A number of studies have demonstrated more transient light responses in parasol cells than in midget cells [39–44]. This difference, previously replicated in large-scale *ex vivo* recordings from macaque retina by observing the more biphasic STA time courses of parasol cells [19,20], was closely mirrored in human RGCs (Figs. 1,4). RF mosaics of the major RGC types in macaque [16,18–20] provided a confirmation of their identity without anatomical measurements, and were broadly similar to the mosaics observed in physiological [26] and anatomical [45] measurements in other species. Similar mosaics were observed in the human RGC types (Fig. 2,8). The precise interlocking of fine RF structure in each mosaic (Fig. 2) also resembled results in macaque [31]. Systematic ON-OFF asymmetries in RF size (Fig. 5) were first observed in macaque retina [16], and examined in greater detail in theoretical work and empirical work in other species [46,47]. Spatial nonlinearities have long been observed in the responses of macaque parasol cells [48– 50], resembling results from cat retina [51,52]. This feature of parasol cells, previously replicated using *ex vivo* recordings [22,32,34] and extended with a computational model that reveals subunit spatial organization and ON-OFF asymmetries [22,34], was also observed in the human data (Fig. 6). Electrical images obtained with high-density arrays (Figs. 3,9) revealed recordings from the somas, dendrites and axons of RGCs, along with the distinctive features of polyaxonal amacrine cells, as was observed previously in macaque [15,23,32,53]. High-resolution visual stimulation previously revealed the structure of RFs in macaque RGCs at single-cone resolution [17,54–57], including the irregular spatial organization of cone inputs seen in human RGCs (Fig. 7). The unusual RF structure of other cell types (Fig. 8,9) has also been examined in macaques, notably in the smooth monostratified cells [20,32,58] and polyaxonal amacrine cells [23,32,53].

The present findings in human retina confirm, and substantially extend, the results of two recent studies, but also differ significantly from the results of a third recent study.

The first recent study [12] functionally classified recorded human RGCs into 5 primary types based on recorded light responses, although the specific cell type identities were not determined and no further analysis of their properties was performed. The results are broadly consistent with the cell type distribution in macaque retina: although there are roughly 20 anatomically distinct macaque RGC types [30], 4-7 of those cell types dominate numerically in physiological recordings [20], and the 5 identified cell types in the human study [12] may have corresponded to ON and OFF midget and parasol cells and the small bistratified cells. In the present work, it is clear that in addition to the ON and OFF midget and parasol cells, which were identified on the basis of density and light response properties, several more types were observed in large numbers, potentially including small bistratified cells, ON and OFF smooth monostratified cells, and polyaxonal amacrine cells, all of which have been studied using the same techniques used here [20,23]. Analysis of additional human cell types (Fig. 2) will be the subject of future work.

The second recent study [13] described the temporal properties and one-dimensional spatial profiles of individual putative parasol and midget RGCs, tentatively identified based on similarities to published work in macaque retina. Although cell type identity was not confirmed, classification was clear, and the results were generally consistent with the present findings. A striking and consistent difference, however, was that the putative human ON-parasol cells exhibited much more biphasic response time courses, with nearly balanced positive and negative lobes, than published data in macaque [16] and the present findings (Figs. 1,2,4), while the other three major types more closely resembled their macaque counterparts. In the present work, only subtle response time course differences were visible between human and macaque RGCs recorded in the same conditions, and these differences were similar in ON-parasol and OFF-parasol cells (Fig. 4). The origin of the discrepancy between the present and preceding study is unclear. One possibility is differences in the visual stimulus parameters. The higher effective contrast of the stimuli used in the previous study -- a field of flickering extended bars rather than flickering small square pixels -- could produce more transient light responses [59– 62]. The bar stimulus also engages more of the RF surround, potentially altering measurements of response dynamics relative to analysis of pixels in the RF center. Because response dynamics and surrounds are mediated by cell type specific circuitry, these effects could be cell type specific. In the present work, the same visual stimulus was delivered to the human retina and to many macaque retinas using the same display, and the same analysis was performed, providing a more direct comparison. Interestingly, in the present data, visual stimulation using larger pixels (not shown) produced more biphasic light responses, particularly in ON-parasol cells, highlighting the importance of matched stimulus conditions. Of course, details of the enucleation procedure or dissection could also affect the results. In sum, it seems likely that the differences between human and macaque ON-parasol cell response time courses in the preceding study are primarily a result of different experimental conditions rather than inter-species differences, but further tests may be valuable.

The third recent study [14] described responses of human RGCs to full-field stimuli with temporal frequency sweeps, full-field contrast steps, modulating gratings, and moving bars. A wide range of responses to these stimuli was observed, but quantitative cell type classification was not performed. This study concluded that the human retina has a substantially greater diversity of functional cell types than macaque retina, and identified only ∼5% of recorded cells that resembled parasol and midget cells, in striking contrast to the numerical dominance of midget and parasol cells in the present results and those of the other recent studies of human retina [12,13], as well as anatomical findings [8,28]. The unique conclusions of this recent study [14] may arise from technical factors. Cell type identification was not confirmed by quantitative clustering (Fig. 2), mosaic organization (Fig. 2) or anatomical techniques. The various stimuli used, although advantageous in some respects, have the disadvantage of simultaneously modulating the activity of many cells, which can produce widespread spike sorting errors that masquerade as variation in response properties across the recorded population. These stimuli also likely recruit wide-field inhibitory circuits, which may be more sensitive to the health of the preparation and tissue dissection. Finally, the human retinas were subjected to prolonged ischemia (7-17m) before isolating the retina, potentially introducing additional variability across cells. Although there certainly may be some cell types in the human retina that are not present in the macaque retina, the convergence of evidence supports the numerical dominance of ON and OFF midget cells, ON and OFF parasol cells, and a few other cell types in the human retina (Fig. 2), as in macaque, and strong quantitative similarities between at least the midget cells and parasol cells in the two species.

Several caveats of the present work bear mention. First, the two recordings analyzed were obtained from a single human retina, though the systematic trends observed in many cells, and the many similarities to previous work in macaque retina, suggest that the findings will generalize. Second, some results were equivocal, for example the well-documented ON-OFF asymmetries in light response dynamics and the contrast-response relationship in macaques [16] were inconsistent in the present data. Third, some properties of the major RGC types have not been confirmed, such as the striking color opponency of the small bistratified RGCs [63,64], the specific S cone inputs to OFF-midget RGCs [17], the synchronized firing properties of major RGC types [65,66], and the distinctive responses to natural scene stimulation [67]. These and other comparisons to macaque retina will await further work.

The present findings support the idea that high-resolution technologies for treating vision loss by stimulation of RGCs (see [68]) can be developed and tested effectively in the macaque retina [69,70], because the numerically dominant RGCs are so similar to those of humans. However, to fully exploit these similarities for the development of future clinical implants will require that the major human RGC types be stimulated with cell type specificity and spatiotemporal precision [71–73]. A companion study will directly probe this possibility.

## Methods

### Retinas

A human eye was obtained from a brain dead donor (29 year-old Hispanic male) through Donor Network West (San Ramon, CA). Macaque eyes were obtained from terminally anesthetized animals used by other researchers in accordance with institutional guidelines for the care and use of animals. Immediately after enucleation, the anterior portion of the eye and vitreous were removed in room light, and the eye cup was placed in a bicarbonate-buffered Ames’ solution (Sigma, St. Louis, MO).

### Multi-electrode array recordings

A custom multi-electrode recording system [15] was used to obtain recordings from human and macaque RGCs in the isolated retina, using methods previously developed and described for macaque recordings [16–18]. In dim light, segments of retina roughly 3 mm in diameter were placed RGC side down on a planar array consisting of 512 extracellular microelectrodes covering a 1.8 mm × 0.9 mm region (roughly 4×8° visual field angle). The primary human retina recording (all figures) was 12mm from the fovea on the superior vertical meridian; this recording was performed with the retinal pigment epithelium attached [17]. A second recording (Fig. 5) was 14mm from the fovea in the superior nasal quadrant; this recording was obtained with the retinal pigment epithelium removed [16,18]. For the macaque retinas, recordings were obtained over eccentricities ranging from 4.5 to 15 mm (4.5 to 12.5 mm temporal equivalent ([16]). For the duration of the recording, the preparation was perfused with Ames’ solution (30-34° C, pH 7.4) bubbled with 95% O2, 5% CO2. The raw voltage traces recorded on each electrode were bandpass filtered, amplified, and digitized at 20kHz [15]. Spikes from individual neurons were identified and segregated by standard spike sorting techniques [15,74,75]. Electrical images were computed for each cell by averaging the voltage waveforms in a time window from 1 ms before to 2 ms after each spike for every electrode on the electrode array.

### Visual stimulation

The visual stimulus was produced by a gamma-corrected CRT monitor (Sony Trinitron Multiscan E100) refreshing at 120 Hz [17], or by a gamma-corrected OLED microdisplay (eMagin) refreshing at 60 Hz [54]. The displays were optically reduced and projected through the mostly-transparent array onto the retina. The light levels at the retina were low photopic: approximately 2200, 2200, and 900 photoisomerizations per cone per second for the L, M and S cones respectively for the CRT display, and 9400, 9300, and 2300 for the OLED display. A spatiotemporal white noise stimulus lasting 30-120 minutes was used to characterize RGC responses [21]. The stimulus consisted of a grid of pixels 2.9-90 µm on a side, updating at 30 or 60 Hz. At each update, the intensities for each of the three monitor primaries at each pixel location were chosen randomly from a binary distribution. The modulation of the three display primaries was either independent or coordinated.

### Cell type classification

The spike-triggered average (STA) stimulus for each neuron was computed from the response to the white noise stimulus [21], to reveal the spatial, temporal, and chromatic properties of light responses. Cell type identification was performed by identifying distinct clusters in a space representing properties of the cells, including the spatial extent and time course, and the spike train autocorrelation (Fig. 2) [16,18,19,26]. In addition, for each neuron, an electrical image (the average spatiotemporal voltage pattern recorded across the array during a spike) was calculated and used for cell type identification and classification [15,32]. This analysis revealed multiple identifiable and complete cell type populations. In particular, the four major types, ON and OFF parasol and midget cells, were readily identifiable by their temporal properties, RF size, density, and mosaic organization (see [16,19,20] for details). The cell densities for human and macaque data were estimated by calculating the typical neighbor distance, and converting to density assuming hexagonal packing. The typical neighbor distance was defined as the median across cells of the median distance to the four nearest, same-type neighbors. Outlier cells, defined as cells with neighbor distance more than three median absolute deviations from the median across cells, were removed prior to calculating the typical neighbor distance. Outlier cells were typically at the edge of the recording or near a gap in the mosaic. These density values were compared to the densities of the four major RGC types expected based on anatomical studies [8,28]. In addition, a number of low-density cell types were identified, including putative small bistratified cells, smooth monostratified cells, and amacrine cells [19,20,23].

### Time course analysis

Time courses for comparison to macaque data (Fig. 4A) were computed by summing the display primary values within the RF from each STA frame, normalizing (L2), and averaging across all cells of each type. The result was then fitted by a cubic spline and upsampled fourfold. For comparison across all cells (Fig. 4B), principal components analysis was performed on the assembled time courses from all preparations, for each cell type. Time courses were then projected onto the first 3 principal components, and a two-dimensional embedding was learned from these projections by applying t-Distributed Stochastic Neighbor Embedding [27].

### Subunit analysis

Subunits were fitted using a method previously described [22]. Briefly, responses to a 15 min white noise stimulus were discretized in 8.33 ms time bins, then divided into training (80%), validation (10%) and testing (10%) partitions. The subunit filters were assumed to be separable in space and time, with the temporal filter identical across subunits and estimated from the STA. Hence, the subunit fitting algorithm only estimated spatial non-linearities.

## Acknowledgements

The human eye was provided by Donor Network West (San Ramon, CA). We are thankful for the cooperation of Donor Network West and all of the organ and tissue donors and their families, for giving the gift of life and the gift of knowledge, by their generous donations. We thank T. Moore, S. Morairty, and the California National Primate Research Center for access to macaque retinas. We thank C. Rhoades for analysis suggestions, and R. Samarakoon, S. Kachiguine and J. Wonderbar for technical assistance. Support: Pew Charitable Trust Scholarship in the Biomedical Sciences (AS), NSF IGERT 0801700 and NSF GRFP DGE-114747 (NB), a donation from John Chen (AML), Research to Prevent Blindness Stein Innovation Award, Wu Tsai Neurosciences Institute Big Ideas, NIH NEI R01-EY021271, NIH NEI R01-EY029247, and NIH NEI P30-EY019005 (EJC).

## Contributions

AK performed dissections and collected the data, AK and EJC conceived and designed the experiments, AK, ARG, NPS, EGW, NB and EJC analyzed the data, AS and AML developed and supported the recording hardware and software, RAS performed the surgery, EJC supervised the project and wrote the paper, all authors edited the paper.

## Notes

### Competing Interest Statement

The authors have declared no competing interest.

### Summary of Updates

Corrected obvious error in abscissa labeling in Fig. 5. No other changes.

